# Rapid QC-MS – Interactive Dashboard for Synchronous Mass Spectrometry Data Acquisition Quality Control

**DOI:** 10.1101/2024.01.30.578059

**Authors:** Wasim Sandhu, Ira J. Gray, Sarah Lin, Joshua E. Elias, Brian C. DeFelice

## Abstract

Consistently collecting high-quality liquid chromatography-coupled tandem mass spectrometry (LC-MS/MS) data is a time-consuming hurdle for untargeted workflows. Analytical controls such as internal and biological standards are commonly included in high throughput workflows, helping researchers recognize low integrity specimens regardless of their biological source. However, evaluating these standards as data are collected has remained a considerable bottleneck – in both person-hours and accuracy. Here we present Rapid QC-MS, an automated, interactive dashboard for assessing LC-MS/MS data quality. Minutes after a new data file is written, a browser-viewable dashboard is updated with quality control results spanning multiple performance dimensions such as instrument sensitivity, in-run retention time shifts, and mass accuracy drift. Rapid QC-MS provides interactive visualizations that help users recognize acute deviations in these performance metrics, as well as gradual drifts over periods of hours, days, months, or years. Rapid QC-MS is open-source, simple to install, and highly configurable. By integrating open-source python libraries and widely used MS analysis software, it can adapt to any LC-MS/MS workflow. Rapid QC-MS runs locally and offers optional remote quality control by syncing with Google Drive. Furthermore, Rapid QC-MS can operate in a semi-autonomous fashion, alerting users to specimens with potentially poor analytical integrity via frequently used messaging applications. Rapid QC-MS offers a fast, straightforward approach to help users collect high-quality untargeted LC-MS/MS data by eliminating many of the most time-consuming steps in manual data curation. Download for free: https://github.com/czbiohub-sf/Rapid-QC-MS

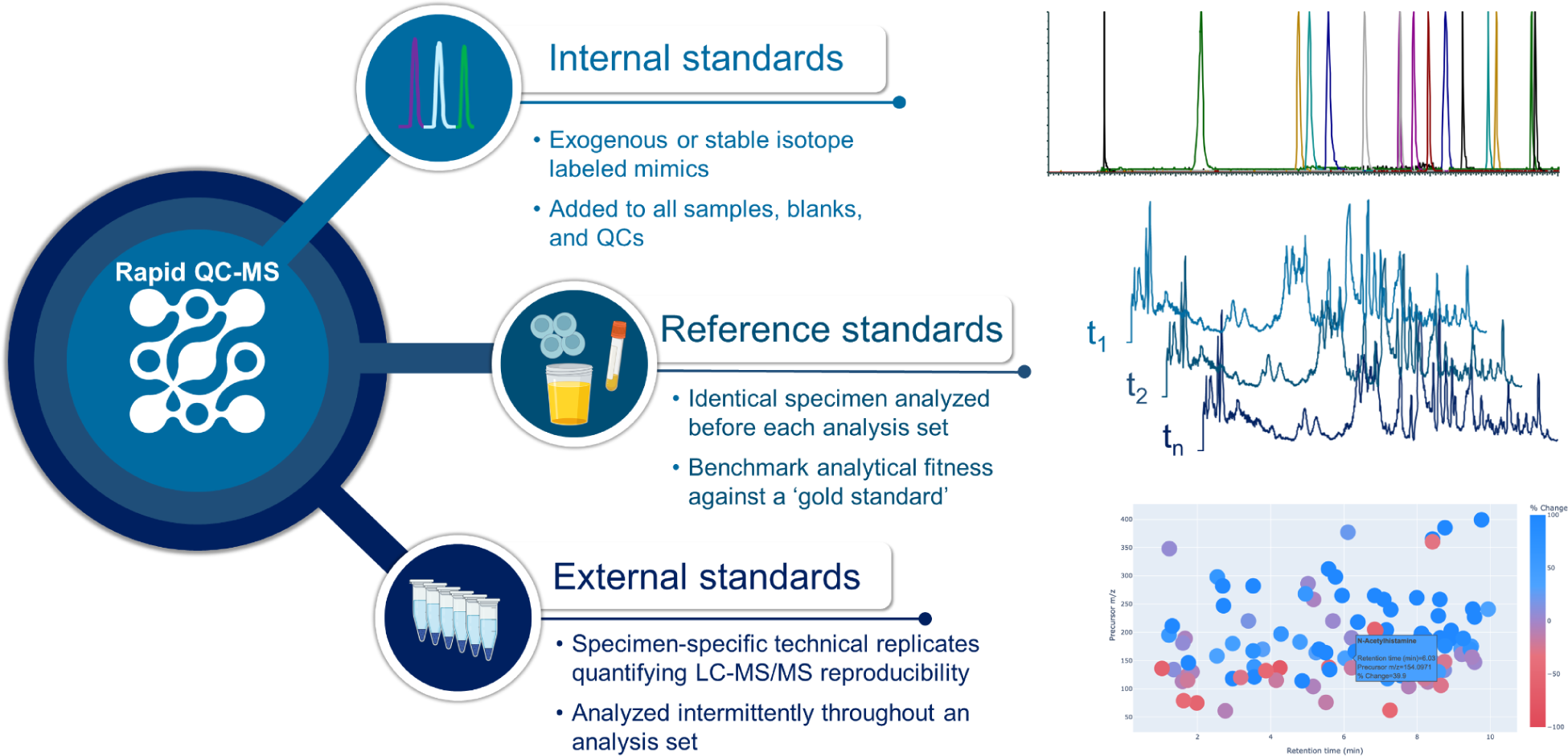

## Introduction

Liquid chromatography-coupled tandem mass spectrometry (LC-MS/MS) is a powerful tool for quantifying biologically relevant molecules such as metabolites, lipids, and proteins. The information generated from these experiments enables deeper understanding of health and disease. Increasingly, automation systems are helping researchers create LC-MS/MS datasets of greater depths spanning thousands to tens of thousands of biological specimens(Cho et al. 2022; Johnson et al. 2019; Li et al. 2020). However, to draw meaningful biological insights, it is critically important that data are of measurably high quality and reproducible. To our knowledge, no freely available software can automatically assess data quality continuously during an analysis. To address this need, we present Rapid QC-MS, software enabling high dimensional LC-MS/MS data quality assessment from anywhere with an internet connection.

Ensuring high quality LC-MS/MS data collection requires monitoring multiple aspects of an analysis(Begou et al. 2018; Dudzik et al. 2018). However, the inherent variability in specimen composition and abundance makes it difficult to compare LC-MS/MS data quality based solely on endogenous molecules. Well-designed experiments should yield multiple quality metrics that assess: sample preparation and injection quality(Simader et al. 2015), instrument performance prior to beginning a new analysis(Lippa et al. 2022), and instrument stability throughout the analysis(Calderón-Santiago et al. 2017). Measuring each of these is essential to creating high quality and high confidence datasets; doing so with minimal delay following data acquisition minimizes sample loss and inefficient use of instrument time.

Internal standards (IS) are exogenous compounds, including stable isotope mimics of endogenous compounds, that are spiked into all specimens, blanks, and controls. By manually extracting IS-specific mass-to-charge ratios (*m/z*), retention times (RT), and intensity information from unprocessed LC-MS/MS data files (raw files), multiple specimens’ fitness can be directly assessed and compared(Evans et al. 2020). Raw files with no detectable IS inform researchers the data collected from the corresponding specimen is of poor analytical integrity. Adding IS at multiple steps during specimen preparation affords greater granularity toward differentiating between extraction and injection effects when troubleshooting the source of poor data quality(Cajka and Fiehn 2014; Broadhurst et al. 2018). IS are versatile: while primarily intended as specimen specific controls, they can also alert users to systemic issues such as shifts in RT and degrading mass accuracy. Continued inconsistency or degradation of signal across multiple specimens can indicate a larger problem with the LC-MS/MS that requires immediate attention to prevent or minimize loss of precious biological material and unproductive instrument time.

Reference standards (RS) typically are complex biological material repeatedly sampled over large periods of time and at least once per analysis(Phinney et al. 2013; Briscoe, Stiles, and Hage 2007). They offer insight into the current state of the LC-MS/MS with respect to its historical performance and to other instruments. Since RS generate many more data points per analysis than IS, they can give more comprehensive insight into an instrument’s analytical capability. Routinely analyzing the same material over the course of weeks, months, and years can reveal changes in several critical metrics including total number of quantified target molecules, target intensities, chromatography, and tandem mass spectra collection. As defined above, RS workflows are analysis and specimen agnostic. They should be evaluated prior to each new analysis set.

External standards (ES) are typically complex reference material used for a specific analysis. They are analyzed periodically over the course of the analysis to more precisely measure LC-MS/MS performance compared to RS. Creating ES by pooling portions of the same biological material being assayed over the course of an analysis allows the researcher to develop QC metrics that are tailored to the molecules being studied. ES workflows, like RS workflows, monitor total number of molecular targets, target intensities. However, unlike RS, ES workflows help evaluate whether specimens analyzed at multiple points during an analysis all produce similar results.

Evaluating IS, RS and ES together (**Figure 1**) provides the user with evidence that any biological conclusions they might draw from their experimental data are not artifacts of poor analytical technique. However, the three types of controls outlined above have a major weakness: their interpretation is usually performed manually, in a time-consuming and subjective fashion. Several computational tools have been described that can assess one or more aspects of the QC workflow outlined above. RawHummus(Dong et al. 2022), an R based shiny app, can be used to assess chromatography and spectral collection across samples and summarize instrument log files. RawHummus could be tailored to investigate IS, RS, and ES but without the granularity of annotated endogenous or exogenous molecules and the added hurdle of manual QC report generation. Other notable tools offer visualization(Avtonomov, Raskind, and Nesvizhskii 2016), IS comparison(Simader et al. 2015), instrument-based bias normalization(Calderón-Santiago et al. 2017) but lack the critical automated nearly instant aspect needed for making rapid data acquisition decisions.

**Figure 1).**
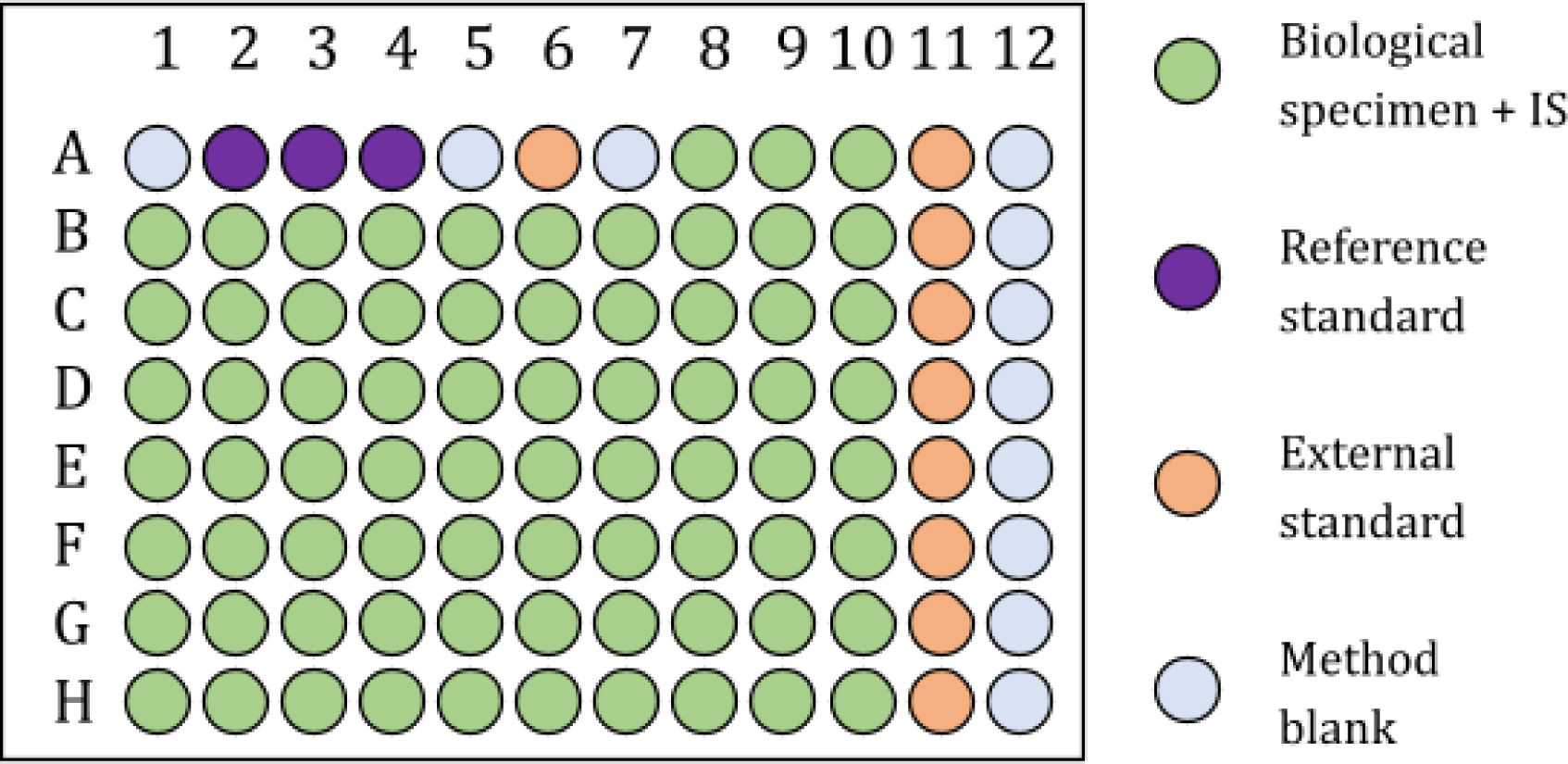
Incorporation of Internal Standards (IS), External Standards (ES), and Reference Standards (RS) in an untargeted workflow. Order of analysis from A1→A12, B1→B12…Order of biological specimens randomized prior to analysis.

In contrast, the Rapid QC-MS software presented here automatically performs rapid quality control over the course of large LC-MS/MS studies. It reports LC-MS/MS quality metrics once each new raw file has been collected and sends an alert to users whenever a newly acquired raw file does not meet specified requirements. Processed mass spectrometry data are stored in a centralized relational database so that specimen-specific QC metrics can be visualized with a browser-based dashboard. This dashboard facilitates intuitive, direct evaluation and comparisons at the specimen (IS), inter-analysis (RS), and intra-analysis (ES) levels. Rapid QC-MS is straight-forward to deploy, scalable across multiple instruments and large specimen numbers, and modularly designed to support community development and new feature incorporation.

## Experimental Section

### Installation and setup

Rapid QC-MS was developed in Python, incorporating several powerful libraries and frameworks frequently used in computational biology: Dash(Hossain 2019), Plotly(*Plotly.py: The Interactive Graphing Library for Python This Project Now Includes Plotly Express!* n.d.), Bootstrap(*Bootstrap: The Most Popular HTML, CSS, and JavaScript Framework for Developing Responsive, Mobile First Projects on the Web* n.d.), Pandas(McKinney 2010; The pandas development team 2023), SQLAlchemy(Bayer n.d.), Watchdog(Mangalapilly n.d.), and the Python APIs for Google Drive and Slack. Rapid QC-MS installation is quick and straightforward with a command line program that configures all components and most dependencies.

However, two dependent software packages, MSConvert (Kessner et al. 2008) and MS-Dial (Tsugawa et al. 2015), (Tsugawa et al. 2020) must be downloaded and installed separately. A full Rapid QC-MS installation guide, including instructions for installing in a virtual environment, is included in the software repository: https://github.com/czbiohub-sf/Rapid-QC-MS/Rapid QC-MS is deployed via command line prompt which launches the user interface dashboard on the local computer’s default web browser. When initially deployed, Rapid QC-MS prompts the user to create a new workspace where the manufacturer of the mass spectrometer is selected. Rapid QC-MS uses manufacturer information to correctly import lists of LC-MS/MS raw files and associated metadata (“acquisition sequences”) from manufacturer-specific formats. Once a workspace has been created, the user can set up synchronous monitoring of IS, RS, and ES by configuring *MS-Dial Configurations, Chromatography Methods*, *Biological Standards and IS Configurations* (Figure 2 and **SI Figure S1**).

**Figure 2).**
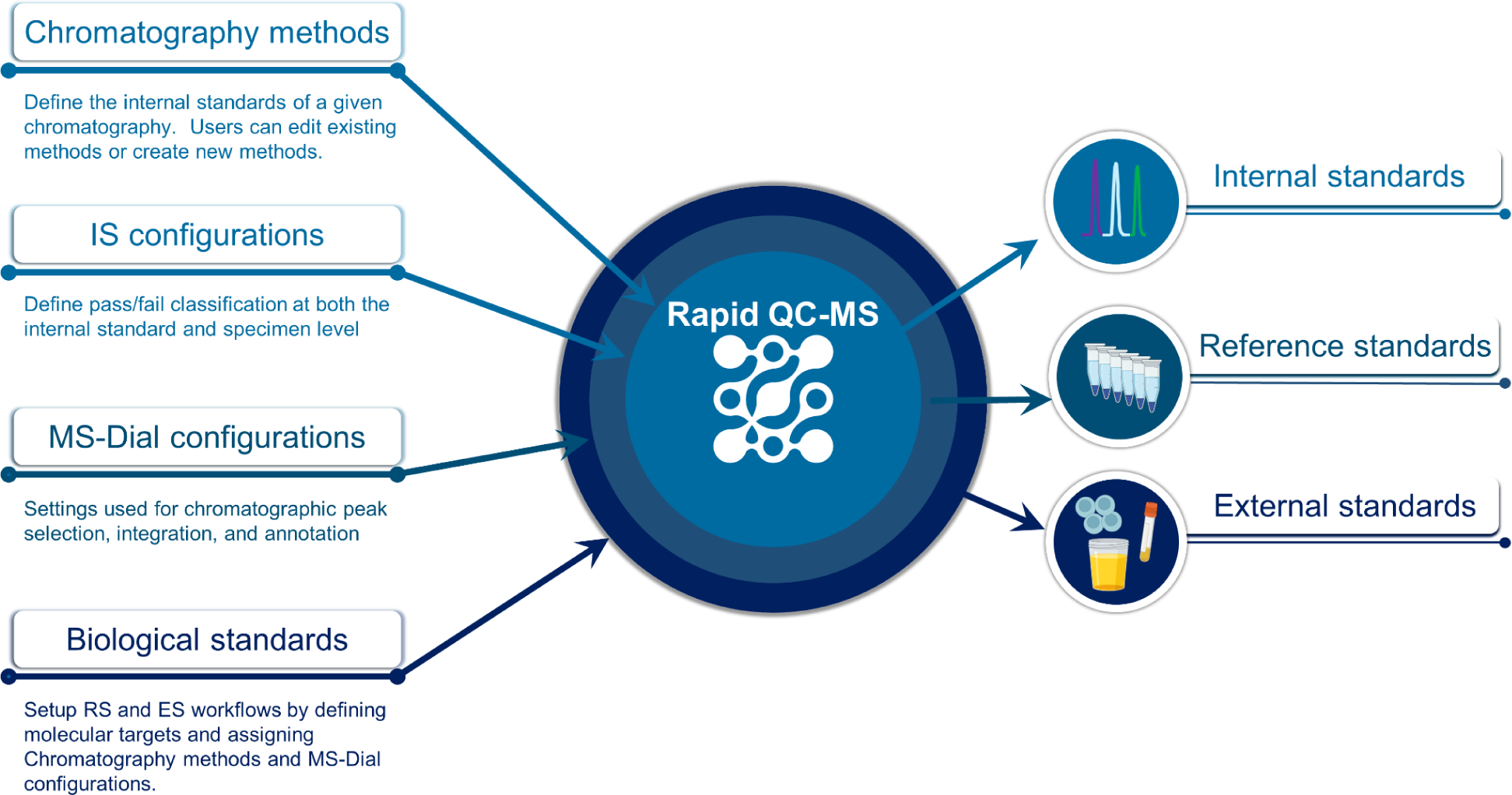
There are four modules that must be set up prior to using Rapid QC-MS. These settings (Chromatography methods, IS configurations, MS-Dial configurations, and Biological standards) dictate what standards Rapid QC-MS searches for, how to quantify and annotate them, and thresholds that must be met to consider a specimen acceptable.

*Chromatography Methods,* applied to IS, RS, and ES workflows, should be assigned for each unique LC-MS/MS method used. Users will provide a unique name for each method, its ionization polarity (positive or negative) and a list of specific IS targets. IS targets must be uploaded for each chromatography method and each polarity (positive or negative modes). Users can upload IS target info as a .msp-formatted file(Kind et al. 2013; Lam and Aebersold 2010) containing IS names, mass-to-charge ratios (*m/z)*, retention times (RT), and tandem mass spectral information (MS^2^). Users who do not wish to use MS^2^ information for IS annotation can upload IS target info as a .csv file containing names, *m/z*, and RT. There is no limit to the number of *Chromatography Methods* that can be created in a workspace.

*MS-Dial Configurations*, applied to all workflows, determine how the MS-Dial backend defines MS1 chromatographic peaks and how those peaks are annotated. Critical parameters include mass accuracy tolerances, minimum peak width and height, and annotation tolerances for RT, *m/z*, and if desired MS^2^ spectral similarity. *MS-Dial Configurations* can be tailored to specific modules. Multiple configurations can be created and saved. Once the user has set up an *MS-Dial Configuration,* previously created *Chromatography Methods* and *Biological Standards* can be updated to include an *MS-Dial Configuration*.

*Biological standards* settings are used to define which specimens and endogenous molecules to extract for carrying out RS and ES workflows. Each biological standard must be assigned a unique text identifier to be extracted from the LC-MS/MS raw filename(s). Rapid QC-MS uses these unique text identifiers to determine which corresponding workflow to apply to each newly generated LC-MS/MS raw file. For example, if a user were to enter the unique text identifier “UrineQC”, every raw file generated with “UrineQC” in the filename would be directed to the predefined “UrineQC” *MS-DIAL Configuration* and annotation library. As with IS targets, users upload an .msp file containing names, *m/z*, RT and MS^2^ spectra of the endogenous targets within RS or ES they wish to track with Rapid QC-MS. The user must also assign *MS-Dial Configurations* and *Chromatography Methods* to these workflows with the user interface.

Creating separate biological standard methods for RS and ES workflows will give users flexibility in how they can be deployed over the course of an analysis set.

*IS Configurations* are exclusively applied to IS workflows. These settings define IS failure criteria for a given LC-MS/MS analysis including the allowable number of missing IS targets, maximum allowable RT shifts, and mass error tolerance. *IS Configurations* are individually named and there is no limit to the number that can be created. Customized *MS-Dial Configurations* can be applied to any *Chromatography Method. IS Configurations* are assigned when setting up a new analysis.

### Design and workflow

Deploying Rapid QC-MS launches the dashboard on the local computer’s default web browser (**SI Figure S2**). The dashboard can be used to initiate new analyses, or ‘Jobs’, review data from completed Jobs, and define settings for IS, RS, and ES workflows. To prepare for a new Job, *Chromatography Methods, MS-Dial Configurations, Biological Standards, and IS Configurations,* as described above, must first be defined. Prior to initiating a new Job in Rapid QC-MS, users must generate an acquisition sequence in the LC-MS/MS system’s control software. Acquisition sequence information can be exported from the vendor software in an open format, typically a .csv file. To initiate a new Job in Rapid QC-MS, the user must click the ‘Setup New QC Job’ button on the dashboard. The user will be prompted to assign a unique Job ID, upload the acquisition sequence, and specify the folder Rapid QC-MS will monitor for newly created raw files (**SI Figure S3**). Once the inputs have been validated, the Job can be started and Rapid QC-MS will monitor newly created raw files with the workflow depicted in Figure 3.

**Figure 3).**
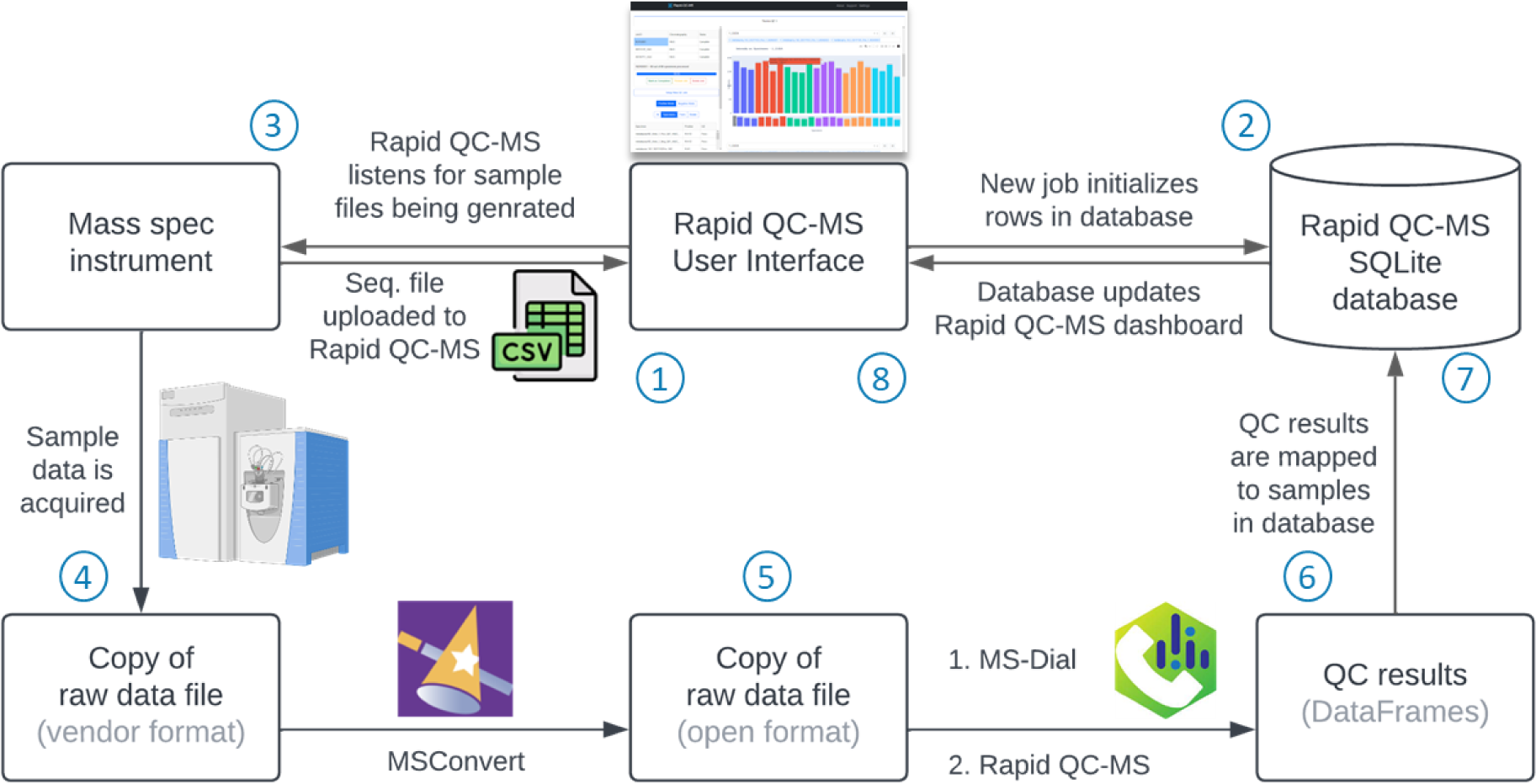
Rapid QC-MS backend for IS, RS, and ES workflows. 1) User initiates new Job and uploads sequence information exported from LC-MS/MS and uploaded to Rapid QC-MS dashboard. 2) New Job initiates new rows in SQLite db, Rapid QC-MS is now listening for new raw files to be created. 3) User initiates analysis in mass spec control software. 4) Once a new file is complete a copy is made to preserve the original file. 5) MSConvert is used to convert from closed vendor format to open format (.mzML) file. 6) MS-Dial searches for and quantifies IS, RS, and/or ES molecular targets. 7) Results are mapped to SQLite db created in step 2. 8) Changes to SQLite database trigger the Rapid QC-MS dashboard to update.

When a new Job is initiated, Rapid QC-MS creates a SQLite database that links the Job with run-specific measurements it will generate. While a Job is active, Rapid QC-MS monitors the previously designated folder for the creation, writing, and completion of raw files. Active monitoring continues until all raw files specified in an acquisition sequence have been generated by the LC-MS/MS and processed by Rapid QC-MS or the user manually marks the Job as complete. Completed raw files are automatically converted to an open format (mzML) via the MSConvert tool(Kessner et al. 2008). The mzML file is then processed by MS-DIAL(Tsugawa et al. 2015), (Tsugawa et al. 2020) which carries out quantification and annotation procedures for all user-defined targets.

When assessing IS, Rapid QC-MS determines pass/flag/fail for each IS target based on parameters defined in the *IS Configurations* settings. Target peak information including intensity, RT, *m/z*, and IS pass/flag/fail status are integrated into Rapid QC-MS’s SQLite database. Any change to the SQLite database triggers the dashboard to refresh, thereby enabling nearly instant review of new QC results. If IS for a given specimen does not meet criteria defined in *IS Configurations*, a failure notification is sent via any supported communication mode to all designated researchers. Rapid QC-MS aligns RS and ES annotations from the current analysis to previously annotated RS and ES specimens respectively. RS and ES specimens are each searched against distinct predefined libraries.

### Cloud sync and notifications

To facilitate instrument monitoring from devices external to the instrument computer, Rapid QC-MS provides secure cloud sync capabilities via Google Drive. By navigating to *Settings* > *General* and entering the required credentials, the user can authenticate and link their Google account with Rapid QC-MS (**SI Figure S4**). The Rapid QC-MS workspace -including SQLite database and .msp/.txt/.csv files - will then be automatically synced to Google Drive storage.

This allows the user to sign into their Rapid QC-MS workspace using their Google account from any device. In the *General* tab users can also register email addresses and business messaging applications to receive Rapid QC-MS notifications of failed or flagged (warning) specimens. For user convenience, these notifications can be turned on or off at any time. Full documentation can be found here: https://czbiohub-sf.github.io/Rapid-QC-MS/user-guide.html

## Results and Discussion

### Rapid assessment of Internal Standards from every specimen

Rapid QC-MS streamlines quality control of active LC-MS/MS analyses by automatically integrating IS peaks and determining if the LC-MS/MS system’s performance is within user defined margins. Poor specimen integrity - measured by shifting RT, drifting *m/z*, or missing IS - automatically triggers notification emails and app-based messages to users. Rapid QC-MS enables nearly instant specimen review through its synchronous interactive browser-based dashboard (Figure 4a-f). Dashboard users select the Job of interest from the Job browser (**4a**). Once a Job is selected, all specimens from that Job populate the Specimen Table (**4e**). Users have the option to filter the displayed LC-MS/MS analyses by their attributes, including ionization polarity or specimen type (**4c**). Within minutes of any new raw file being completed, the user can efficiently monitor RT, intensity, and Δ*m/z* data from all IS (**Figure 4b,d,f**). Intensity, RT, and Δ*m/z* plots are malleable and can be dynamically modified and filtered by IS or specimen(s) using auto-filled dropdowns. Furthermore, by supplying additional experiment-specific metadata from a supported .csv file, Rapid QC-MS automatically groups and color codes specimens according to any treatment/class information provided **(4d)**. This utility can help the user determine if any observed IS variance is due to matrix effects from a specimen treatment, a systematic error, or an unknown stochastic process. An example metadata file can be found in **Supporting Information**

**Figure 4).**
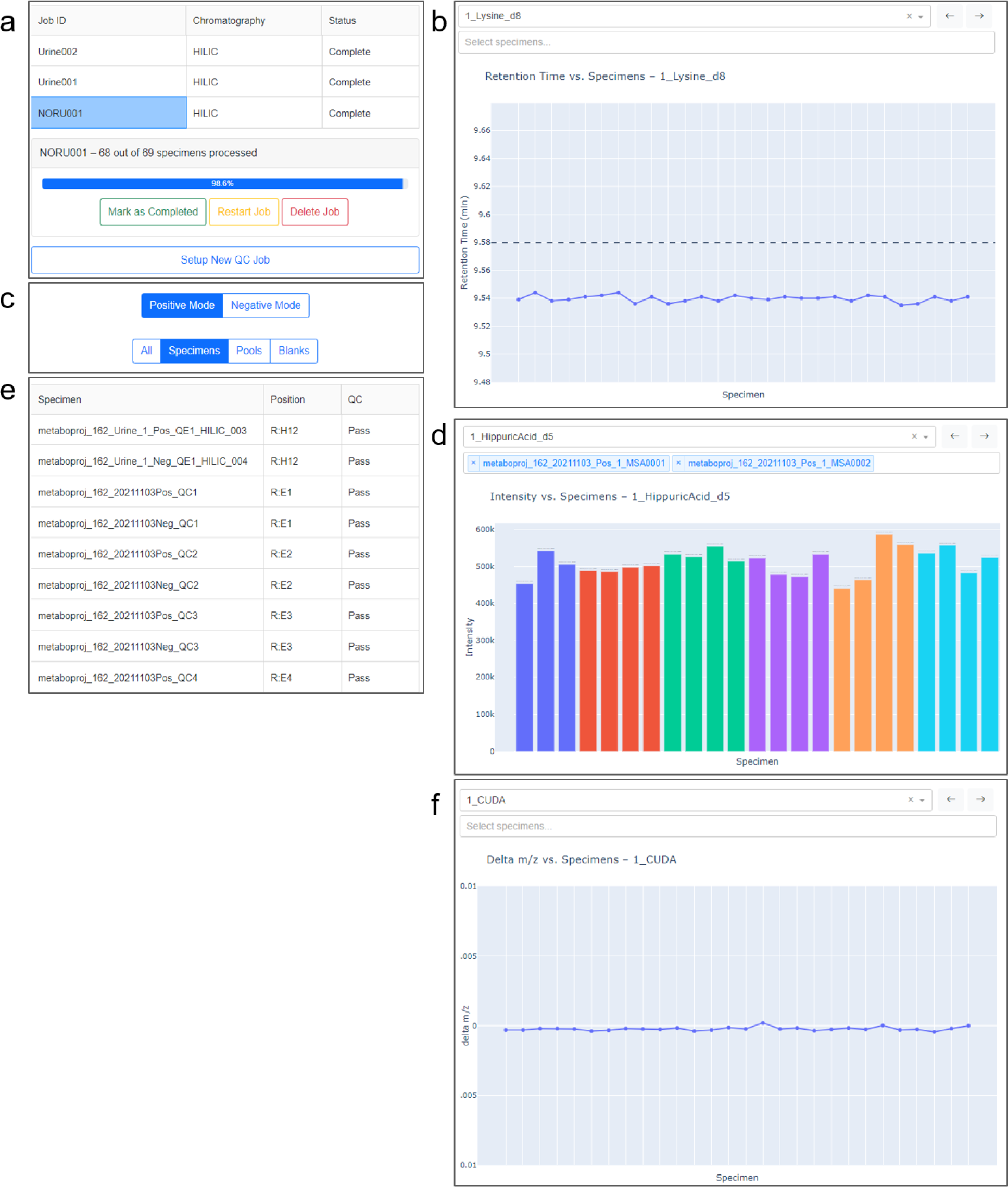
The Rapid QC-MS dashboard. **a)** Jobs overview; Lists past and present jobs. New jobs can be created via the ‘Setup New QC Job’ button. **b)** IS RT across specimens, in order of analysis. **c)** Control panel to filter visualized data by ionization polarity and type. **d)** IS intensity plots across specimens, colors denote specimen class as defined in the optional metadata table. **e)** List of all specimens within a Job that have already been processed with Rapid QC-MS **f)** IS *m/z* error across specimens.

The status of all specimens in a Job can be found in the dashboard’s Specimen Table (**Figure 4e**). The QC column indicates the status, ‘Pass,’ ‘Fail,’ or ‘Warn’, of each specimen that has passed through the pipeline. The thresholds for QC pass and fail are explicit, defined by the user (*Settings* > *IS Configurations*). A “Warning” label alerts users to specimens that narrowly pass QC, and are assigned if either of the following conditions are met: The number of IS intensity dropouts is 75% or more of the defined threshold; or 50% or more IS have a QC “Warning” designation as described below.

Clicking on a filename in the Specimen Table will load the corresponding Specimen Information Card (**SI Figure S5**). The Specimen Information Card contains the QC results of each IS target within a selected specimen, which are also assigned ‘Pass,’ ‘Fail,’ and ‘Warn’ designations as follows: Missing IS are marked ‘Fail’. Rapid QC-MS assigns ‘Warning’ to IS with Δ*m/z* or RT or outside the pass criteria. “Warnings” for individual IS could be triggered if any of the following are true: 1) The ΔRT is greater than 66% of the “RT shift from library value” threshold; 2) The in-run ΔRT is greater than 80% of the “RT shift from in-run average value” threshold; 3) The Δm/z is greater than 80% of the “RT shift from library value” threshold.

### Instrument benchmarking with a Reference Standard

Rapid QC-MS simplifies the use of RS for benchmarking instrument performance and system suitability testing. Integrated into the Rapid QC-MS dashboard, the RS workflow offers users a quick assessment of the current state of the LC-MS/MS instrument compared to previous analyses. The RS workflow is flexible and can be performed multiple times in a Job, once per Job, or excluded all together. Applicable to any specimen type, RS assesses a user-defined list of molecular targets based on *m/z*, RT, and (optional) MS^2^ matching to highlight changes in sensitivity across hundreds or thousands of molecular targets. The RS workflow gives users the ability to compare the current state of the LC-MS/MS system to a running historical average of each molecular target or a single, pre-designated “gold standard” raw file (**Figure 5**).

**Figure 5).**
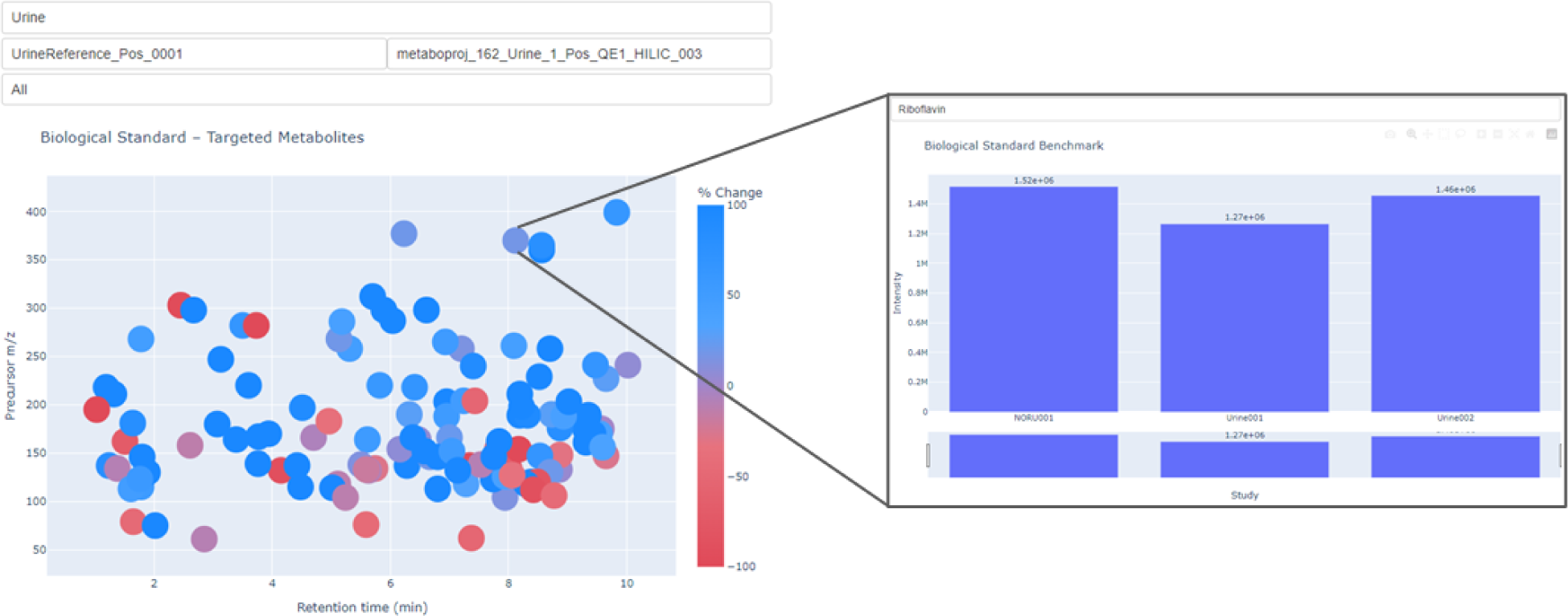
Biological standards viewing module from Rapid QC-MS dashboard. Users can view all annotated features from an RS or ES specimen and compare intensities to previously analyzed specimens from within a job or across jobs. Users can select any feature for additional information including annotated name, RT and *m/z*.

To create a benchmarking workflow in Rapid QC-MS, a user first generates a list of high confidence annotations in a well-defined biological matrix. Next, they combine name, *m/z*, RT, and MS^2^ information in a .msp-format file which they then upload into Rapid QC-MS by navigating to *Settings* > *Biological Standards.* Selecting endogenous analytes with fragment-rich MS^2^ spectra and minimal MS^1^ signal interference will ensure the most accurate peak integrations by the MS-Dial backend.

### External Standards workflow

Biological standards are versatile and can be applied to control multiple aspects of an analysis. Setting up an ES workflow is nearly identical to building an RS workflow, however Rapid QC-MS separates the two making it easier to simultaneously evaluate the scope of each analysis with greater granularity. The key distinction is the molecular targets library used to annotate ES has a much broader scope compared to the highly tailored RS library. Because the composition of ES often varies between analyses an all-encompassing library is recommended for ES workflows. We suggest creating an .msp file containing only molecular targets with known RT corresponding to the LC-MS/MS method being employed. Not all molecular targets will be detected in a single specimen matrix; but a broad library allows for a baseline of hundreds or thousands of molecular targets, a subset of which will be quantified in any biological specimen. Example raw files, .msp files, metadata and sequence files can be can be found in the **Supporting Information** or on the Rapid QC-MS website: https://czbiohub-sf.github.io/Rapid-QC-MS

### Modular design with an emphasis on documentation

Rapid QC-MS is fully open-source, built modularly with thorough documentation, and is designed to be expanded by its users and integrated into existing experimental and automation workflows. The software was developed to be easy to install and use, scalable to large amounts of data, and flexible enough to support the development of new features. It also offers a range of features for data management and organization, including secure cloud synchronization, nearly instant notifications, and centralized storage in a local database. We have found Rapid QC-MS to be a useful tool for assessing the quality and reliability of mass spectrometry data in untargeted workflows spanning thousands of samples. Its automated quality control capabilities, combined with its data management and visualization features, make it an indispensable tool for researchers.

Together, the generated suite of plots offers a nearly instant and convenient way to verify and quantify RT shifts, *m/z* drifts, and IS intensities during analysis. The RT across specimens plot visualizes deviations from expected RTs, quickly pointing the user to investigate chromatography. The intensity across specimens plot gives insight into injection and ionization efficiency, enabling users to broadly assess specimen quality by comparing the peak heights of IS. The Δ*m/z* across specimens plot is useful for verifying the mass accuracy of the measurements made during the run, ensuring that identifications are reliable for downstream analysis. Additionally, Rapid QC-MS provides researchers with the ability to quantify their instrument’s performance using the Reference Standard workflow. Finally, we offer the External Standard workflow to monitor specimen specific changes to endogenous molecules across the length of an analysis.

## Conclusion

The Rapid QC-MS dashboard offers a fast and effective way to ensure the collection of high-quality untargeted LC-MS/MS data. Rapid QC-MS can be applied with any chromatography or mass spectrometry methods. By automatically evaluating numerous parameters within minutes of writing a specimen’s raw file, Rapid QC-MS allows users to quickly and easily determine the fitness of each specimen and the analysis in total. The dashboard also offers interactive visualizations that allow users to monitor nuanced changes in their data, such as RT shifts, *m/z* drifts, and systemic changes like intensity fluctuations. The Rapid QC-MS package is easy to install, modular, and highly configurable, making it adaptable to a variety of workflows and pipelines. Primarily developed with Thermo Fisher orbitrap-MS, sequence files from other LC-MS/MS systems can be easily appended for use with Rapid QC-MS. Overall, the use of Rapid QC-MS can save time and effort in the quality control process, freeing up researchers to focus on conducting experiments.

## Acknowledgements

We sincerely thank Carlos Gonzalez for software testing and troubleshooting. We thank Jonathan Liu, Samuel D’Souza, and Angela Pisco for input during tool development. We thank the organizers of US-HUPO 2023 and the Metabolomics Association of North America 2023 for allowing us to present this work and the fruitful discussions and suggestions it fostered. We thank Abby Sarkar and Roel Nusse for publicly releasing our collaborative data, which is available at the Metabolomics Workbench (ST002263) repository and can be used with the supporting information as demonstration data. BioRender was used to generate some images in figures 2 and 3. This work is supported by the Chan Zuckerberg Biohub - San Francisco and its donors Priscilla Chan and Mark Zuckerberg.

## Supporting information

**Supporting Figures (S1-S5)** are attached below. These figures illustrate the Rapid QC-MS user interface and add context at useful areas of the manuscript.

Additional supporting information is available for download at our project repository (https://github.com/czbiohub-sf/Rapid-QC-MS) or downloaded directly (https://doi.org/10.5281/zenodo.10525183):

- Example data can be downloaded from the NIH Metabolomics Workbench: http://dx.doi.org/10.21228/M8Z119 ; this data accompanies author provided supplemental materials including sequence files (.csv), metadata file (.csv), and chromatography specific MS2 spectral library files (.msp).
- Six total .raw files containing urine data used as reference standards. Two files accompany Metabolomics Workbench study ST002263, these files were not appropriate to be uploaded with the larger study as they were only used to gauge the health of the LC-MS/MS, not applied within study ST002263
  - The remain four .raw files can be used to demonstrate the reference standard comparison module
- Four .csv files
  - Sequence files for creating new Rapid QC-MS Jobs utilizing the demonstration data provide
  - A metadata file to accompany ST002263
- Six .msp library files.
  - Reference standard, negative mode
  - Reference standard, positive mode
  - External standard, negative mode
  - External standard, positive mode
  - Internal standards, negative mode
  - Internal standards, positive mode

**Supporting Information Figure S1).**
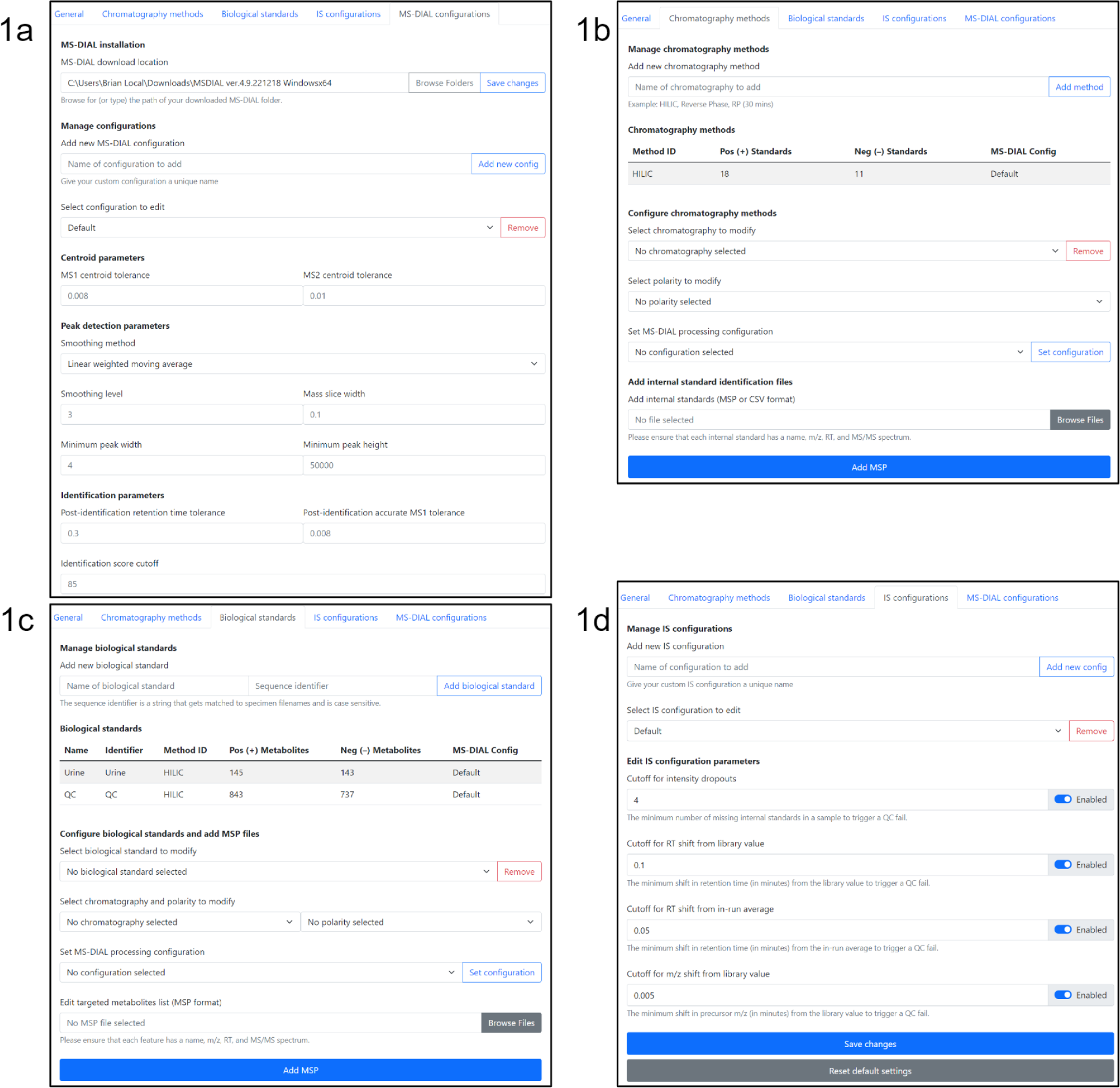
Rapid QC-MS configurations enable the user to define criteria for quantification of IS, ES, and RS features. **1a** *MS-DIAL configurations* tab; users set parameters for peak integration and annotation. **1b** *Chromatography methods* tab is used to define the IS of a given chromatography. Users can edit existing methods or create new methods. **1c** *Biological standards* tab is used to set up RS and ES workflows by defining molecular targets and designating *Chromatography methods* and *MS-Dial configurations*. **1d** *IS configurations* are used define pass/fail classification in IS workflows

**Supporting Information Figure S2).**
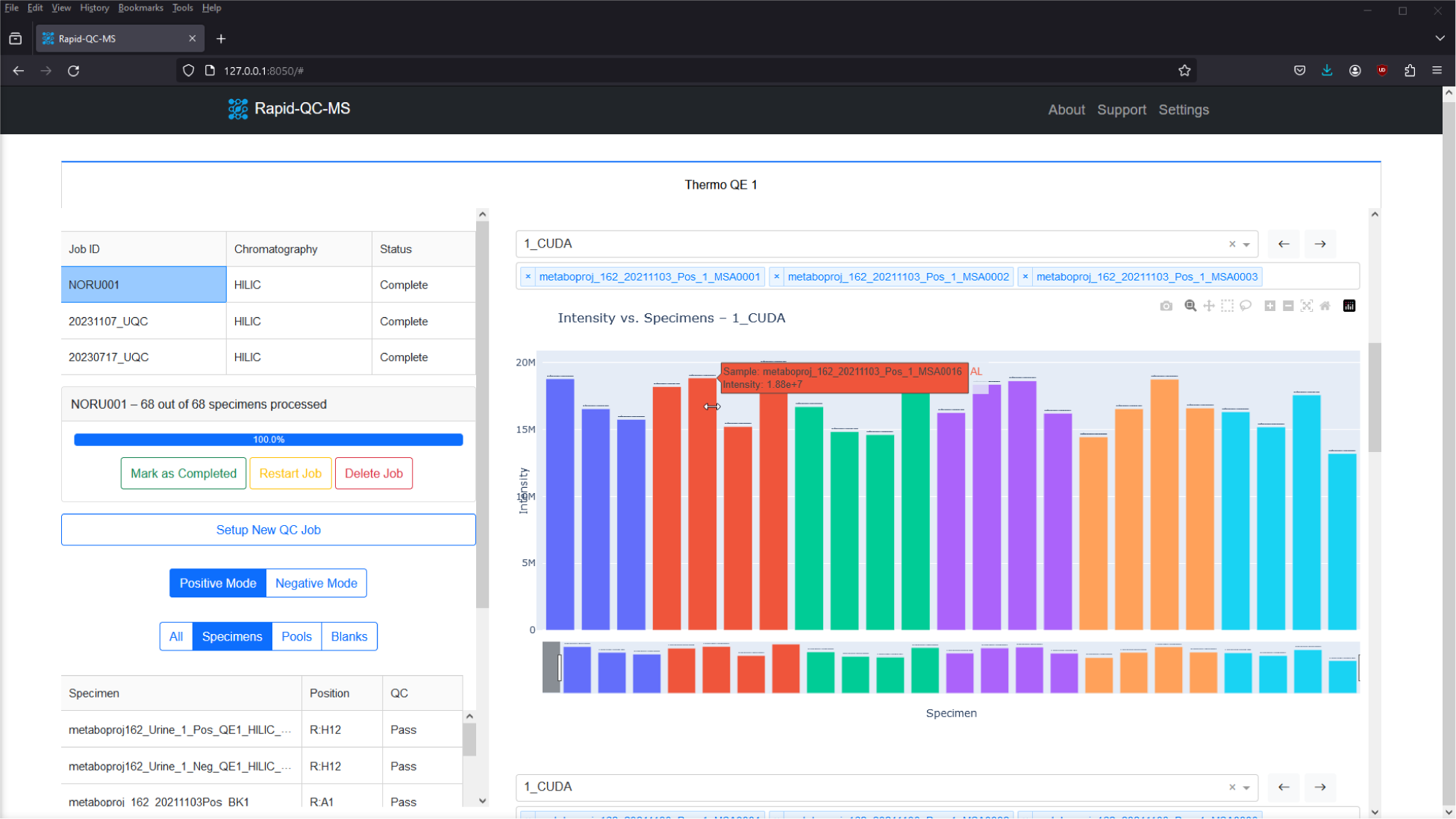
When Rapid QC-MS is deployed it launches the web-browser based dashboard where users can review current and archived Jobs or create a new Job or setup additional methods and configurations.

**Supporting Information Figure S3).**
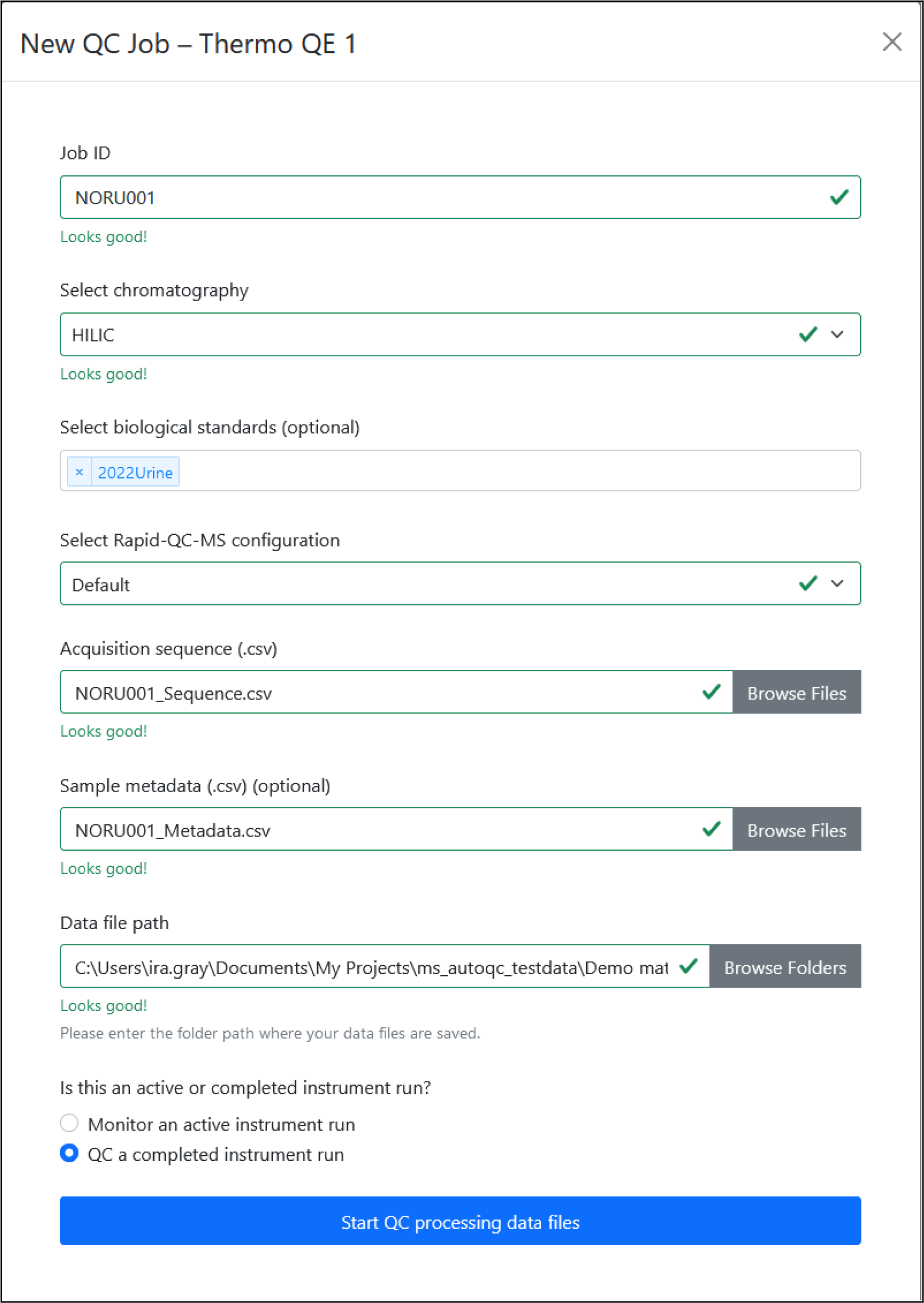
Creating a new Job in Rapid QC-MS. Each Job requires a unique Job ID. Additionally, users must select the relevant *Chromatography method, IS Configuration*, and provide the acquisition sequence. Users also have the option to select biological standards and upload specimen metadata.

**Supporting Information Figure S4).**
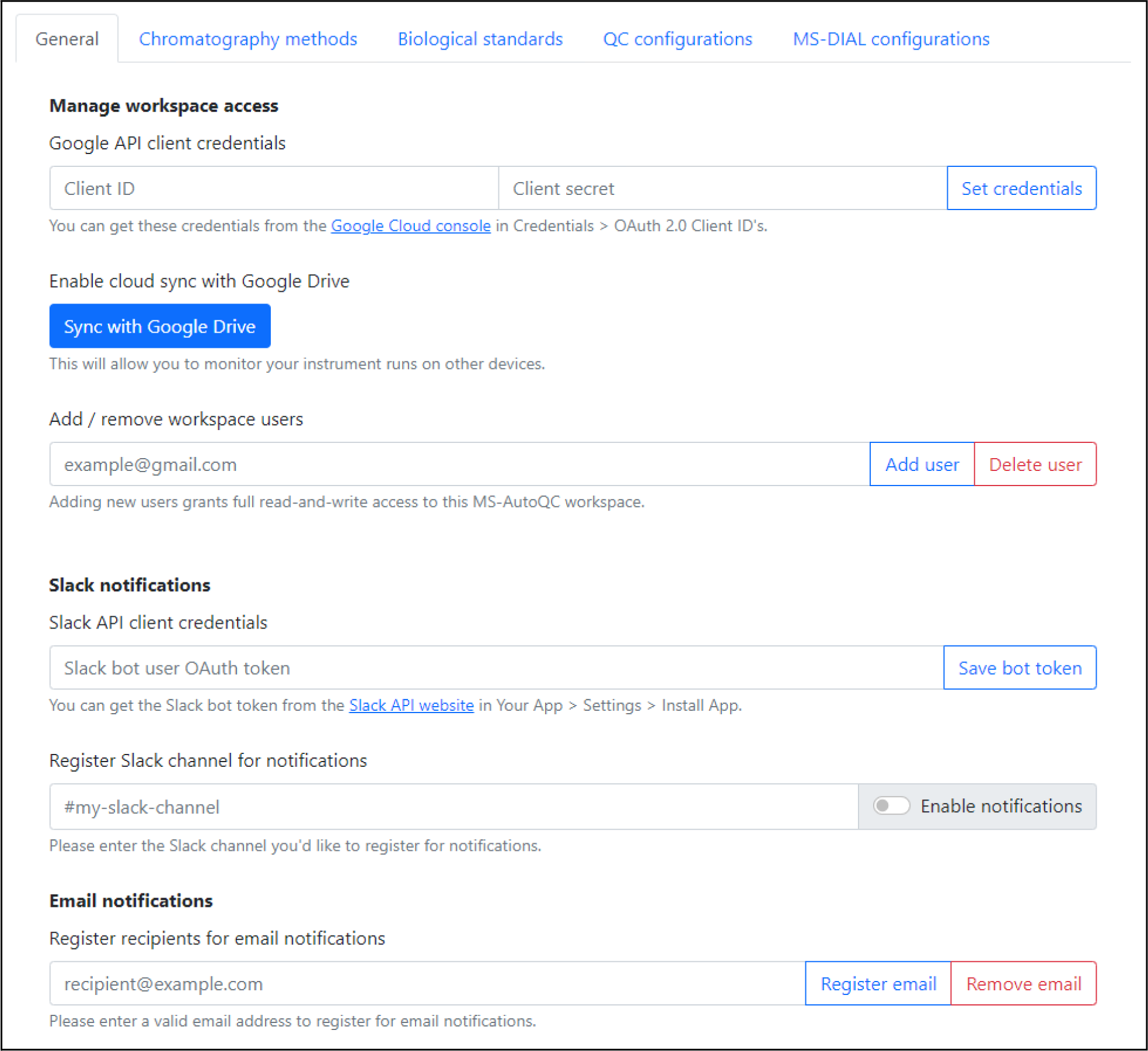
The *General* tab can be used to set up external messaging and dashboard access. Found by navigating *Settings → General* from the Rapid QC-MS dashboard.

**Supporting Information Figure S5).**
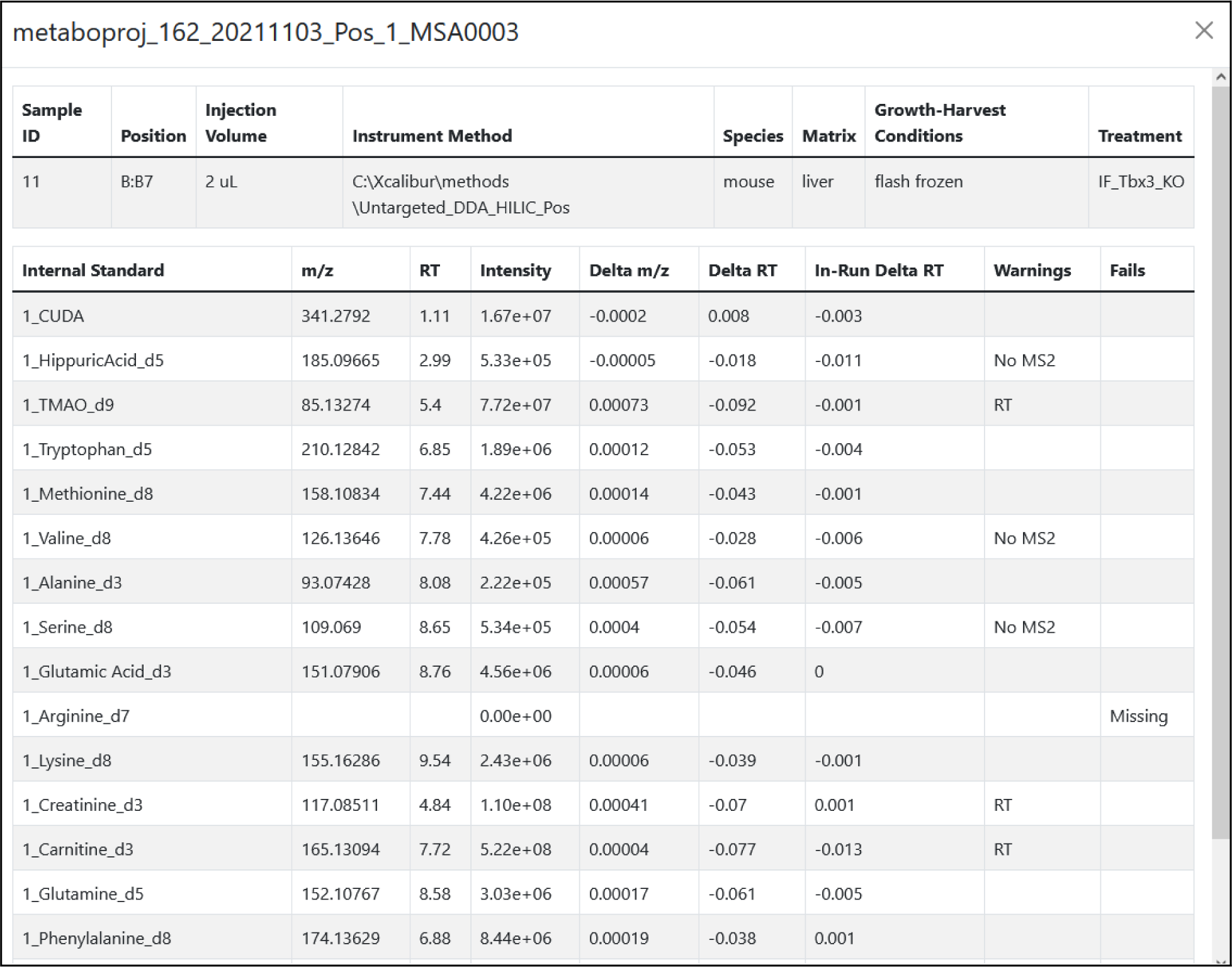
Specimen information card. From the dashboard users can click on a specimen for IS level pass/fail/warn information and in the case of fail and warn provide justification. Also listed are IS intensity, mass error (Delta *m/z*), RT error (Delta RT), and in-run RT shift (In-Run Delta RT).

## Notes

https://github.com/czbiohub-sf/Rapid-QC-MS

https://doi.org/10.5281/zenodo.10525183

http://dx.doi.org/10.21228/M8Z119

### Competing Interest Statement

The authors have declared no competing interest.

### Summary of Updates

Author order was incorrect

